# Being bigger is better: Size-dependent advantages of plasmodial fusion in a multinucleated slime mould

**DOI:** 10.1101/2025.01.31.634296

**Authors:** Cathleen M. E. Broersma, Daniel Rozen

**Affiliations:** Institute of Biology, Leiden University, Leiden, The Netherlands

**Keywords:** size benefits, cell fusion, chimera, social evolution, *Physarum polycephalum*, slime mould

## Abstract

Larger body size often enhances survival and reproduction. While most organisms grow to achieve larger sizes, some can rapidly increase size through fusion. Fusion decisions are influenced by genetic relatedness and environmental factors, balancing potential benefits and costs. In *Physarum polycephalum*, a multinucleated slime mould, fusion is thought to occur mainly between genetically identical or highly related individuals. However, the frequency of fusion among close relatives, and the drivers and benefits of fusion, remain unclear. This study explored fusion frequency, morphology, and benefits among clonal and closely related individuals of *Physarum*. We also assessed the impact of abiotic stress on fusion and whether genetic relatedness between fusion partners influences survival during exposure to stress. Our results revealed variation in fusion frequency and morphology among different plasmodial lines and pairs of close relatives. Abiotic stress increased fusion rates between both clonal and compatible non-clonal pairs. However, the benefits of fusion did not depend on the whether fusion occurred between clones or non-clones. These findings demonstrate plasmodial fusion provides size-related survival benefits. Furthermore, the promotion of fusion in stressful conditions highlights its adaptive role in responding to environmental challenges. Collectively, these results underscore the evolutionary importance of fusion in *Physarum*, shaping its ecological success and resilience.

**Competing Interests Statement:** The authors declare no competing financial interests.

## Introduction

Larger organismal size can be associated with increased survival and reproductive output (Buss 1980; Highsmith, Riggs, and D’Antonio 1980; Hughes and Connell 1987; Harvell and Grosberg 1988; Foster et al. 2002; Bonner 2004, 2006). These fitness benefits may result from enhanced resource acquisition, greater competitiveness in social interactions, and improved defence against predators and environmental stressors (Brooks and Dodson 1965; Paine 1969; Buss 1980; Rinkevich and Weissman 1992; Foster et al. 2002; Smith, Queller, and Strassmann 2014; Kapsetaki and West 2019).

Whereas becoming larger typically results from growth, some organisms have evolved strategies that allow them to increase in size more rapidly through fusion with compatible partners (Buss 1982; Glass, Jacobson, and Shiu 2000; Roca, Read, and Wheals 2005). Fusion is found in diverse taxa, including species of plants, fungi, animals, cyanobacteria, and protists, and is associated with a range of potential benefits (Buss 1982; Stoner, Rinkevich, and Weissman 1999). For example, hyphal fusion between compatible colonies of the fungus *Neurospora crassa* can result in increased spore production (Rayner 1991; Bastiaans, Debets, and Aanen 2015). Similarly, fusion between coral larvae of colonial marine hydrozoan species can increase growth rate and survival via increased filter-feeding capacity and improved structural integrity (Buss 1990; Feldgarden and Yund 1992; Amar, Chadwick, and Rinkevich 2008). Because fusion can enable organisms to quickly form large structures without relying solely on prolonged growth, it allows fused individuals to rapidly gain size-related benefits (Buss 1982; Madelin 1984; Buss 1999).

However, fusion may come with costs. Fusion can allow horizontal transmission of infectious cytoplasmic elements, pathogens and viruses (van Diepeningen, Debets, and Hoekstra 1997; Debets and Griffiths 1998; Aanen et al. 2008; Paoletti and Saupe 2009). Additionally, when genetically different individuals fuse to form a chimera, co-occurrence of nuclei and cytoplasmic elements of both parties creates opportunities for one individual to exploit the other (Trivers 1971; Axelrod and Hamilton 1981; Sachs et al. 2004; Burt and Trivers 2006; Foster and Wenseleers 2006). This selfishness may be expressed as social exploitation of resources or reproductive competition between the cell lines (Buss 1982; Debets and Griffiths 1998). Hence, whether two individuals successfully fuse will depend on whether the potential benefits associated with fusion will outweigh its associated costs (Aanen et al. 2008).

Given the potential costs of fusion, theory predicts that mechanisms should evolve that enable individuals to restrict fusion to desired individuals (Hamilton 1964a, 1964b; Aanen et al. 2008; Bourke 2011). Indeed, all organisms that undergo fusion have evolved specialized allorecognition systems that make it possible to identify conspecifics and distinguish self vs non-self (Grosberg and Quinn 1988; Buss 1990; Crampton and Hurst 1994; Crespi 2001; Wilson and Grosberg 2004). As a result, fusion is generally restricted to clones and, in rare cases, to closely related individuals or kin. Fusion between close kin can be explained by kin-selection theory which posits a decrease in potential costs with increasing relatedness between interactants (Hamilton 1964a; Buss 1982; Buss and Grosberg 1990). At the same time, fusion between closely related (non-clonal) kin may be a consequence of an imperfect self-recognition system rather than dependent on potential benefits of fusion with kin (Feldgarden and Yund 1992; Hughes et al. 2004; Aanen et al. 2008). Regardless, the widespread existence of allorecognition in fusion-capable organisms suggests that on average the potential costs of fusion between non-clonal individuals outweigh its potential benefits (Wilson and Grosberg 2004; Aanen et al. 2008).

The slime mould *Physarum polycephalum* is an ideal organism to investigate fusion between close relatives and explore its potential benefits and costs. In its vegetative stage, *P. polycephalum* forms a giant multinucleated, or syncytial, network structure called a plasmodium, which, despite its large size, is a single giant cell. Importantly, compatible plasmodia can fuse to form a single network, resulting in an almost instantaneous increase in size (see examples in Figure 2). To our knowledge, size-related benefits and costs of fusion have not been empirically studied in *Physarum* but are likely conditional on the abiotic and social (biotic) environment. For instance, larger networks might access spatially distributed resources more effectively, share nutrients, and dilute stressors in adverse environments (Buss 1982; Rayner et al. 1984; Tröger, Goirand, and Alim 2024).

Conversely, when individuals reach a sufficient size or fusion is energetically costly, further fusion may offer diminishing benefits, and an excessively large plasmodium may impair coordinated transport and homeostasis (i.e., network functioning). Additionally, as seen in other fusion-capable organisms, fusion carries the risk of social exploitation or the transmission of harmful cytoplasmic elements. These hypotheses, however, remain untested.

Allorecognition in *P. polycephalum* is regulated by up to sixteen loci that act at different stages of the interaction between plasmodia, both on the cell surface and within the joined cytoplasm (Carlile and Dee 1967; Collins 1972; Schrauwen 1979; Betterley and Collins 1984; Clark and Haskins 2012). Briefly, fusion compatibility (i.e., whether two plasmodia successfully fuse) is assessed at three different stages: before contact, during contact, and upon mixing of the cytoplasm after fusion (summarized in Clark and Haskins 2012).

Successful fusion occurs only when interacting plasmodia are genetically similar at each locus and incompatibilities at any stage lead to rejection or termination of fusion. As a result, a wide range of fusion morphologies and compatibilities can be observed, including successful fusion, temporary fusion, and even lethal reactions post-fusion where one or both parties is negatively impacted (Carlile and Dee 1967; Schrauwen 1979; Clark and Haskins 2012). The complexity of fusion and the genetic diversity of compatibility loci implies that the rates of fusion in nature are extremely low unless strains are clonal or very closely related (Clark and Haskins 2012). Consequently, little is known about drivers and consequences of clonal and non-clonal fusion in*. polycephalum*.

Here, we first investigated whether increased size through growth provides a survival benefit in *P. polycephalum*. We then examined whether fusion occurs between closely related plasmodia, either among progeny or between parent-progeny, and the nature of these interactions. To explore potential non-genetic factors influencing fusion, we tested whether stress exposure affects fusion rates in pairs consisting of clonal and fusion-compatible non-clonal plasmodia. We then assessed if fusion, resulting in increased size, similarly confers a survival benefit. Finally, to look for evidence that fusion among non-clonal plasmodia might promote selfish behaviour and consequently reduces network functioning, we compared the magnitude of the size- benefits between clonal and non-clonal pairs. Our findings reveal a clear survival benefit associated with larger size, achieved through either growth or the rapid size increase following fusion. Moreover, fusion in compatible non-clonal pairs provided survival benefits comparable to those observed in clonal pairs, suggesting no evident costs associated with non-clonality in these cases.

## Materials and Methods

### Species

*Physarum polycephalum* is a plasmodial slime mould found in humid, shaded environments such as the forest floor, in the leaf litter, and on decaying logs (Madelin 1984). In its plasmodial stage it exists as a diploid, multinucleated, single celled organism that feeds on other microbes through phagocytosis (Fig. 1). When environmental conditions become worse, it can form sclerotia which are dormant desiccation-resistant structures. Alternatively, the plasmodium can produce fruiting body structures called sporangia in which haploid spores are formed through meiosis. When conditions are favourable, a spore germinates to release a single amoeba. Two mating compatible amoebae can fuse (i.e., mate) to form a zygote in which nuclear fusion takes place to produce a diploid cell. Through repeated cycles of nuclear division, *without* cell division, the zygote then develops into a multinucleated plasmodium, completing the life cycle. Notably, there are thus two stages in which fusion takes place: nuclear fusion during the mating stage and plasmodial fusion during the vegetative stage—in this study we focus on plasmodial fusion (red dashed area in Figure 1).

**Figure 1.**
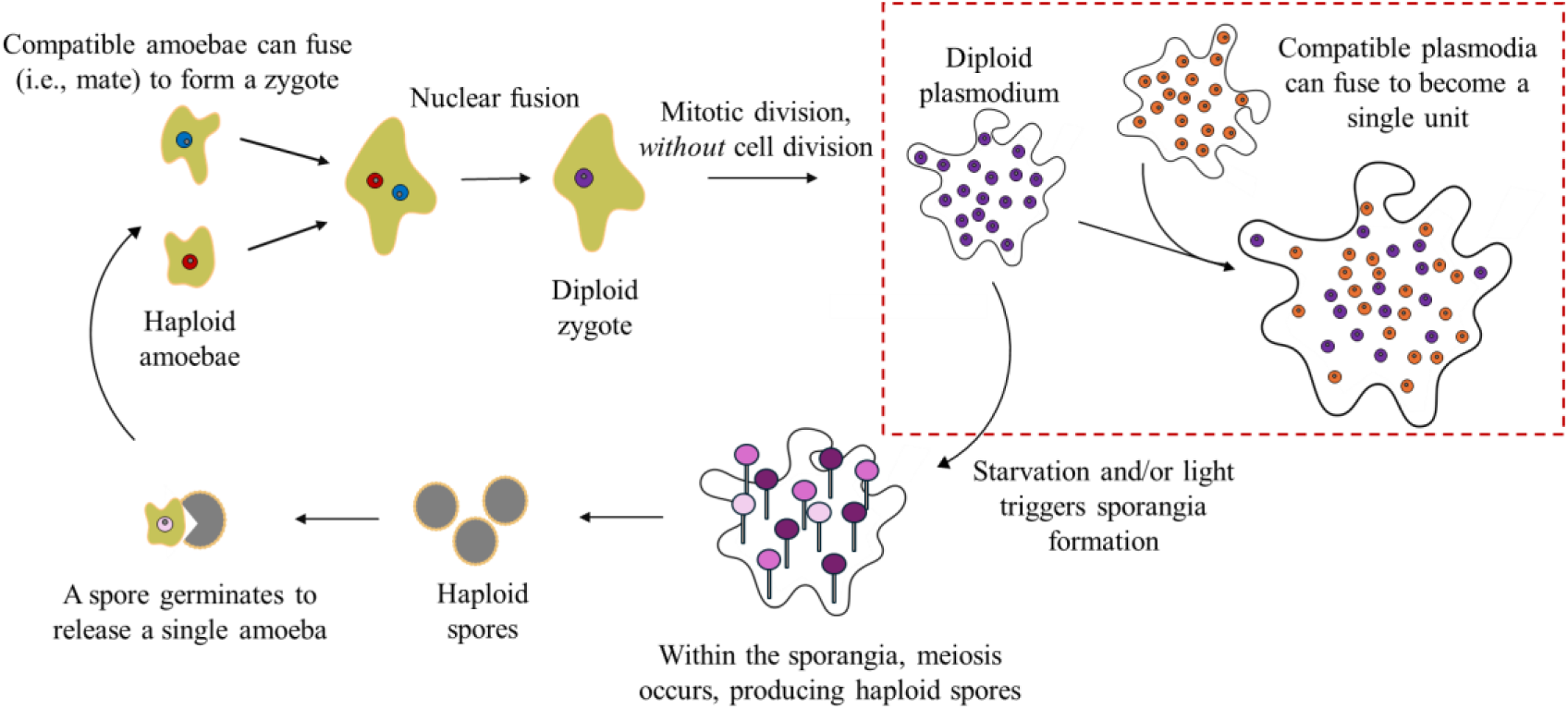
The life cycle of *Physarum polycephalum*. The figures are not drawn to scale. Details are discussed in the text.

**Figure 2.**
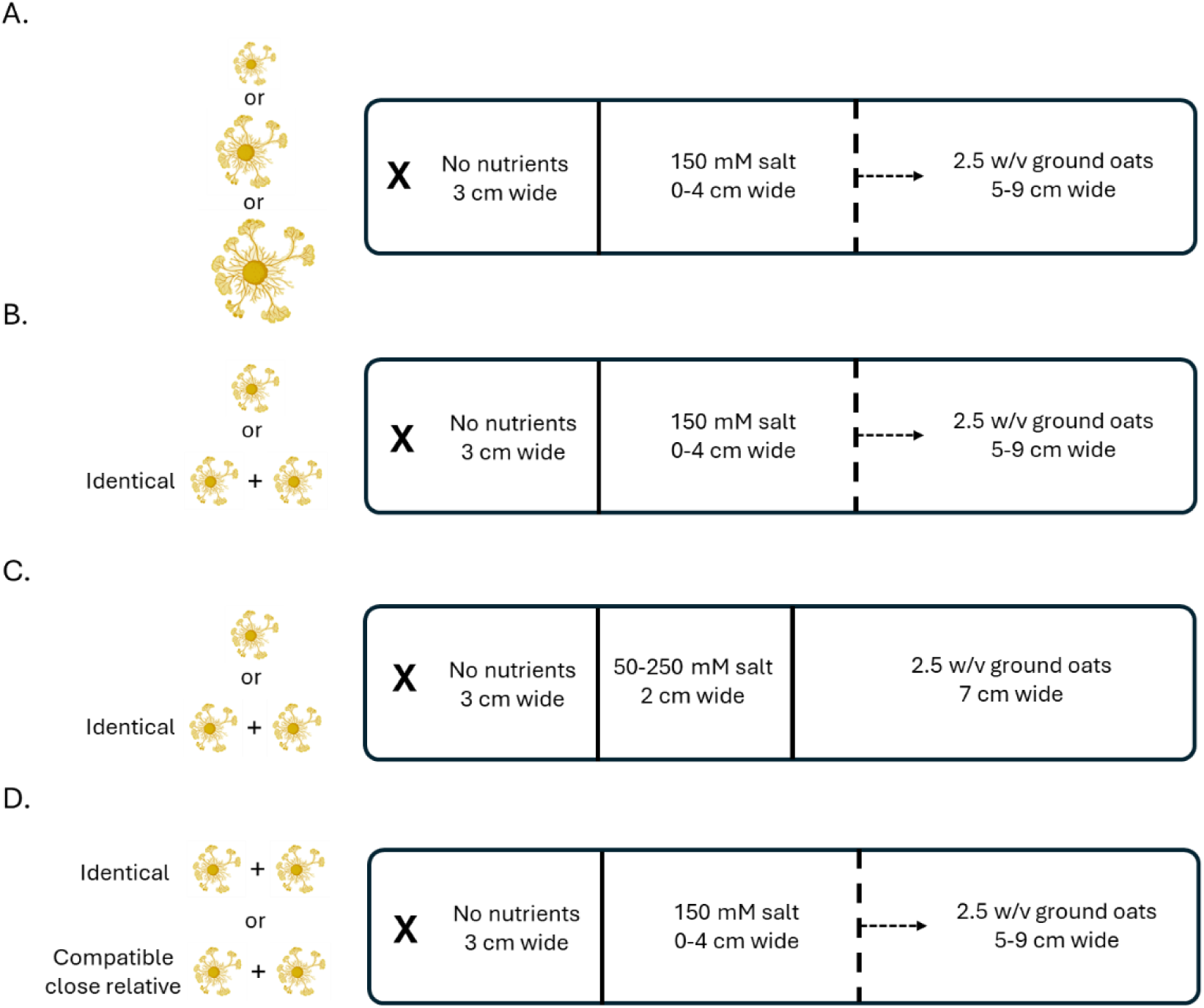
Experimental set-ups. The ‘X’ denotes the point(s) of inoculation of the plasmodia. (A) To examine the effect of size on distance salt compartment crossed, plasmodia of three different sizes were inoculated in the first compartment. (B) To examine the effect of fusion with a genetically identical (clonal) plasmodium on the width of the salt compartment crossed, a single plasmodium or two clonal plasmodia were (co-)inoculated. (C) To examine the effect of fusion with a clonal plasmodium on concentration of the salt compartment crossed, a single plasmodium or two clonal plasmodia were (co-)inoculated. (D) To examine the effect of genetic identity of the fusion partner, i.e., clonal vs compatible close relative (non-clonal), on the width of the salt compartment crossed, two plasmodia were co-inoculated.

### Strain culturing

We used the amoebae of *P. polycephalum* strains AI16, AI35, TU41, DP246, and KM88 obtained from S. Kawano (Professor emeritus at The University of Tokyo). The amoebae of each strain were grown on PGY plates (0.5 % glucose, 0.05 % yeast extract, 2 mM MgSO4, and 2% agar in 25 mM potassium buffer, pH 6.6) at 25°C in the presence of live bacteria (*Klebsiella pneumonia*) as a food source. To induce zygote formation, we harvested the amoebae and washed them once in BSS (3.0 g citric acid monohydrate, 4.2 g K2HPO4•3H2O, 0.25 g NaCl, 0.21 g MgSO4•5H2O, 0.05 g CaCl2•2H2O, pH 5.0) via centrifugation at 400 *x* g for 3 min. We resuspended the amoebae in 200 μl BSS and counted the cell density using a hemacytometer. We then mixed the amoebae of two strains at equal ratios (∼1 x10^6^ amoebae total) in a total of 100 μl, added 200 μl of dense bacterial culture, and spread the mixture on a fresh PGY plate. After 3-5 days, amoebae had fused (i.e., mated) to form zygotes that further developed into individual microplasmodia (Fig. 1). We isolated a single microplasmodium (∼3mm diameter), transferred this to a non-nutrient agar (2%) plate with sterile oat flakes, and allowed the network to cover the plate. From these plates, we prepared stocks of sclerotia that could be revived later. Using this method we established four plasmodial lines, hereafter called “parental” plasmodial lines: AI35.TU41, KM88.AI35, DP246.KM88, and A16.TU41.

As sixteen fusion loci need to be identical (or highly similar) for successful plasmodial fusion in *P. polycephalum*, the chances that two non-clones fuse are extremely small. To obtain plasmodial lines that could potentially fuse, we established closely related lines derived from three of the parental plasmodial lines: AI35.TU41, KM88.AI35, and DP246.KM88. To do so, we induced spore formation by exposing a plate full of network to light for two days at room temperature (on the bench top). We then placed the plate back in the dark at 25°C. Using this method, sporangia containing spores formed within five days. Note that during spore formation, meiosis takes place to produce haploid spores—meaning that the spores are siblings that share roughly 50% of genetic material. We waited another five days for the spores to mature and dry out. We then collected all the spore heads from a single plate using tweezers, transferred a few spore heads to a vial containing sterile tap water, mixed well, and counted the spore density using a hemacytometer. To induce spore germination, we inoculated 1x10^5^ spores in sterile tap water (pH adjusted to 5.5). After five days, we harvested the mixture of spores, cysts (encysted dormant amoebae), and amoebae, centrifuged them, and resuspended them in 300 μl live *K. pneumonia* culture. We spread this mixture onto a PGY plate to allow the amoebae to grow in numbers by division and form zygotes. Daily, we checked for the formation of microplasmodia emerging from the zygotes using a microscope. As soon as microplasmodia became visible (8-11 days after inoculation), we transferred individual plasmodia to water agar plates containing sterilized oat flakes, making sure to transfer a single plasmodium. We did this for a total of fourteen plasmodia, hereafter called “progeny” plasmodial lines, for each of the three parental plasmodial lines.

## Experimental procedures

### Benefits of size

We first examined if plasmodia of different sizes varied in their survival, here tested as their ability to overcome salt stress. Salt is a well-studied repellent known to cause osmotic stress and toxicity in *P. polycephalum* (Vogel and Dussutour 2016). Briefly, we used a 3-compartment setup in plastic containers (11.8 x 2.8 cm, 50 ml polystyrene reservoir, Oxygen, USA) (modified from Nagarajan-Radha and Beekman (2023)). The first compartment was 3 cm wide and contained demi water agar, the centre compartment varied between 1 and 4 cm wide and contained demi water agar with 150 mM NaCl, and the third compartment was 5-8 cm wide and contained demi water agar with finely ground oats (2.5% w/v) (Fig. 2A). All media were prepared with 2 % agar. We inoculated a plug with a part of a growing plasmodial network in the first compartment. The plates were then placed in a large plastic box with a lid (not fully closed) and incubated in the dark at 25°C. After 24 hours, we scored if plasmodia had crossed the salt compartment into the oats compartment. We tested networks of three sizes: 7, 18, and 45 mm^2^. Each plug size was tested thirteen times using different plasmodial lines on different days.

### Testing for fusion compatibility

We tested fusion compatibility among all progeny plasmodial lines from each parental line in all pairwise combinations, and between progeny plasmodia and their parental line. This resulted in a total of 120 pairwise combinations for each parental line (among *N*=14 progeny lines and *N*=1 parental line), which included each line tested against itself. Note that because we allowed mating to take place among the sibling amoebae, a pair made up of progeny plasmodia share roughly 50% of their genetic material. A pair made up of the parent and an offspring plasmodium also share roughly 50% of their genetic material. To monitor the interaction between plasmodia, for each pairwise combination, we placed either a plug from the leading edge of a plasmodium or an oat flake covered with plasmodium in the opposite corners of wells in a 25-well plate well (∼2 cm apart)

containing non-nutrient agar (2 % in demi water). We imaged the wells using a Zeiss Zoom V16 microscope at 80X magnification for 10-16 hours with 30-minute intervals to determine the nature of the fusion interaction. Each plasmodial pair was tested at least three times on different days. From the videos, we classified fusion compatibility into two main groups: *i.* successful fusion and *ii.* rejection. Successful fusion can be observed by the formation of one or multiple major veins that arise shortly after plasmodia make contact (Fig. 3A). Within the category of rejection, we further distinguished between *iii.* overgrowth, where plasmodial grow on top of each other without fusing (Fig. 3B), *iv.* avoidance, where plasmodia after becoming physically close (with or without physical contact) move away from each other (Fig. 3C), *v.* rapid retraction, where one or both plasmodia almost instantly retract their network(s) after physical contact (Fig. 3D), and *vi.* transitory fusion, where plasmodia initially fuse and form one or multiple major veins, but within hours again split (Fig. 3E).

**Figure 3.**
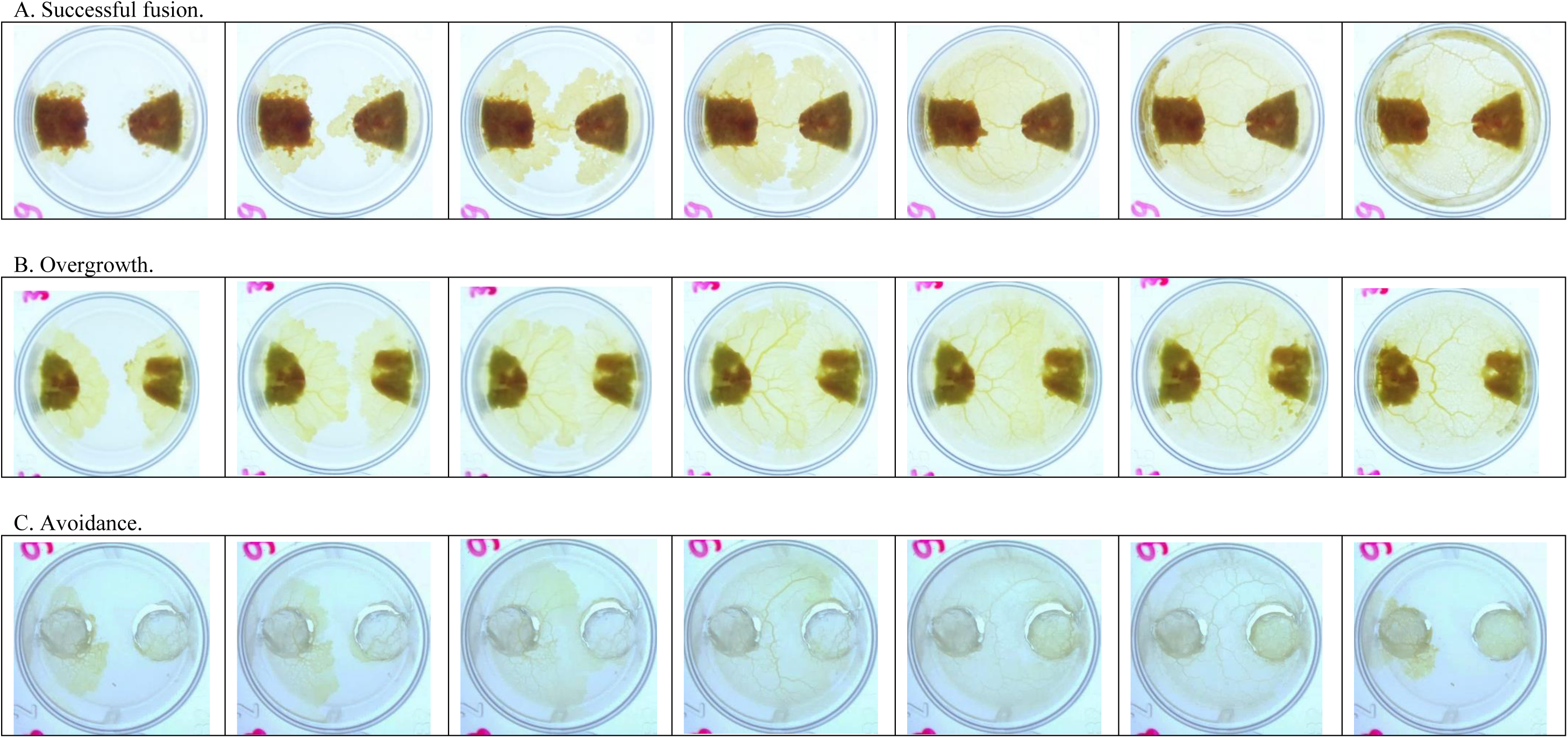
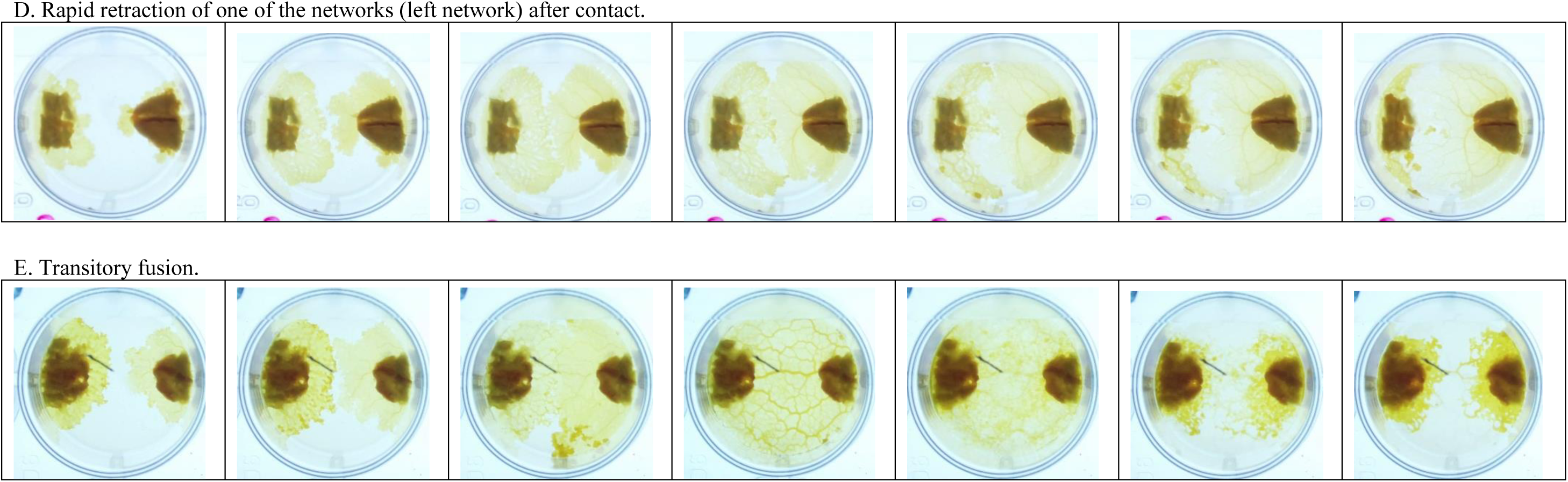
Examples of categories of fusion morphologies in pairs of closely related plasmodia of *Physarum polycephalum*. (A) Successful fusion. (B) Overgrowth. (C) Avoidance. (D) Rapid retraction of network(s). (E) Transitory fusion. Images shown are a subset of those captured across a 24-hour time course with images taken every 30 minutes.

### Testing the effects of the abiotic and biotic environment on fusion rates

We examined whether the rate of fusion was influenced by the genetic identity of the fusion partner (clonal versus fusion-compatible non-clonal partner) and exposure to salt stress. To test this, we assessed fusion in pairs of plasmodia consisting of clonal and non-clonal partners, under conditions with or without a 150 mM salt bridge.

### Benefits of fusion

We examined if fused clonal plasmodia could cross *i.* a wider salt compartment (Fig. 2B) and *ii.* a compartment with a higher salt concentration (Fig. 2C) than individual plasmodia. To test the first hypothesis, we used the same setup as described above, with a few modifications. Specifically, here we inoculated either a single plasmodium or two clonal plasmodia, placed ∼4 mm apart, in the first compartment (Fig. 2B). The size of each plug was 18 mm^2^, i.e., the same size as the middle-sized plug in the first experiment. After 24 hours, we scored if *i.* plasmodia fused before crossing the salt bridge and *ii.* crossed the salt compartment into the oats compartment. We excluded samples in which the plasmodia did not fuse before crossing the salt compartment. We performed these experiments with six plasmodial lines which were replicated at least three times on different days (*N*=3-9).

To examine if fused clonal plasmodial lines could cross a centre compartment with increasing salt concentrations, we used the 3-compartment set-up with the following modifications. The first compartment was 3 cm wide and contained demi water agar, the centre compartment was 2 cm wide, and the oats compartment was 7 cm wide (Fig. 2C). The salt concentration in the centre compartment ranged from 50 to 200 mM with steps of 25 mM (= seven concentrations tested in total). As a control (0 mM), we added demi water agar to the centre compartment. We performed these experiments with four plasmodial lines which were replicated three times on different days.

We next examined if size benefits obtained through fusion depended on social partner, i.e., clonal versus compatible non-clonal fusion partner. To do so, we used the 3-compartment set-up where we varied the width of the centre salt compartment (Fig. 2D). We inoculated either a pair of clonal or compatible non-clonal plasmodia, placed ∼4 mm apart in the first compartment. We performed these experiments with four focal progeny plasmodial lines (offspring of parental line KM88.AI35) and replicated each pairwise combination a total of eight times on two different days.

### Statistical analyses

We performed all analyses in R (version 4.2.1) (R Core Team 2023).

To test for benefits of network size, we modelled the maximum distance of the salt compartment crossed as a function of network size. The model was fit using maximum likelihood with the ‘glmmTMB’ function with a Gaussian error structure. We included network size as a fixed effect and assessed its significance through type II Wald chi-square tests using the Anova function from the ‘car’ package. We included plasmodial line and block as random effects, and assessed their significance through single deletions of terms, comparing the reduced models and full model using a likelihood ratio test. Pairwise comparisons between group means were conducted with adjustments for multiple comparisons using the Tukey HSD method using the ‘emmeans’ package.

To test if the abiotic environment and/or identity of the fusion partner (clonal partner vs a compatible non-clonal) influenced fusion frequency, we modelled fusion as a function of absence/presence of salt and the genetic identity of the fusion partner. The model was fit using maximum likelihood with a beta binomial error structure. We included salt presence (yes vs. no), genetic identity of the fusion partner, and their interaction as fixed effects. We included block as a random effect. The significance of the terms was assessed as described in the first model.

To test whether the benefits of greater size were influenced by fusion, we performed two models to examine: *i.* the maximum distance crossed of the salt compartment, and *ii.* the maximum salt concentration crossed, both as functions of partner presence (no partner vs clonal partner). In both models, we included partner presence as a fixed effect and focal line and block as random effects. The models were fit using maximum likelihood with the ‘glmmTMB’ function with a Gaussian error structure, and the significance of the terms were assessed as described in the first model.

To test if benefits of greater size were influenced by with whom you fuse (clonal partner vs a compatible non-clonal), we modelled the maximum distance of the salt compartment crossed as a function of the genetic identity of the fusion partner. The model was fit using maximum likelihood with the ‘glmmTMB’ function with a Gaussian error structure, and the significance of the terms were assessed as described in the first model.

## Results

### The benefits of being larger on survival

Before testing whether fusion confers size-related benefits, we first assessed the effect of size on survival in plasmodia of *P. polycephalum* in the absence of fusion. Specifically, we examined whether initial plasmodial size influenced an individual’s ability to cross a 150 mM salt bridge of varying widths. We found a significant effect of size on the distance of the salt compartment crossed, where larger networks could cross wider salt compartments (Fig. 4; 𝜒^2^=158.47, *df*=2, *P*<0.001). There was also an effect of plasmodial line (𝜒^2^=8.628, *df*=1, *P*=0.003), indicating that some lines could cross wider salt compartments compared to others.

**Figure 4.**
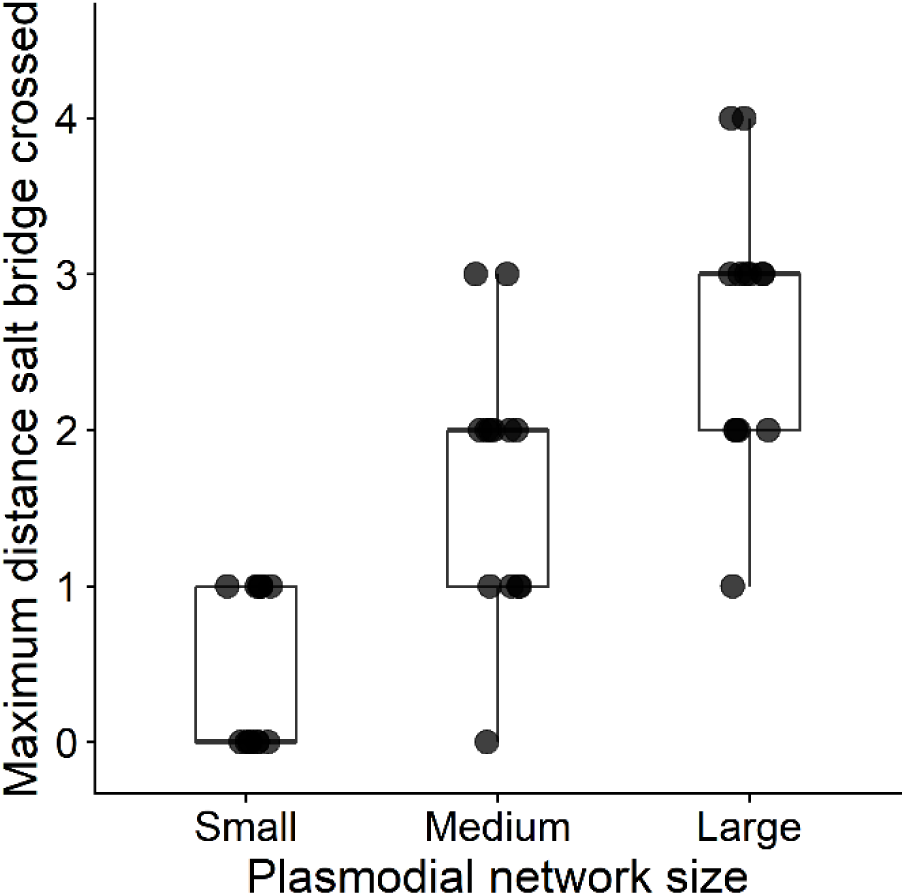
The maximum distance (in cm) of a 150mM salt bridge crossed by plasmodia of different initial sizes. Plasmodial network sizes tested were small (7 mm^2^), medium (18 mm^2^), and large (45 mm^2^), i.e., a ∼2.5-fold increase in plasmodial size with increasing plug sizes.

### Variation in fusion morphology within and between plasmodial lines

To obtain pairs of genetically different plasmodial lines that could fuse, we examined the fusion morphology among pairs of progeny from a single parental line, and between all progeny and their parents (*N*=120 pairs among *N*=14 progeny + 1 parental line). We did this in three different parental lines (AI35.TU41, DP246.KM88, and KM88.AI35). Table 1 shows a summary of the outcomes of the fusion morphology observations. We found diversity in the propensity to fuse, with examples of successful fusion and rejection in each of the three parental lines (Fig. 3). Notably, the frequency of successful fusion varied between parental lines, where fusion was successful in 3% of pairs in AI35.TU41 and 8% in DP246.KM88, in contrast to 63% in pairs of KM88.AI35. Within the category of rejection, we could further distinguish between *i*. overgrowth, where plasmodial grew on top of each other without fusing, *ii*. avoidance, where plasmodia after becoming physically close (with or without physical contact) moved away from each other, *iii*. rapid retraction, where one or both plasmodia almost instantly retracted their network(s) after physical contact, and *iv*. transitory fusion, where plasmodia initially fused and formed one or multiple major veins, but within hours again disconnected to become two networks.

**Table 1.** The observed fusion morphology in pairs of progeny and pairs of progeny with its parental line.

|  |  | AI35.TU41 | DP246.KM88 | KM88.AI35 |
| --- | --- | --- | --- | --- |
| fusion | successful fusion | 4 | 9 | 75 |
| rejection | total | 101 | 96 | 30 |
|  | overgrowth | 87 | 85 | 12 |
|  | avoidance | 0 | 0 | 11 |
|  | rapid retraction of one or both networks | 10 | 0 | 7 |
|  | transitory fusion | 4 | 11 | 0 |

### The influence of the abiotic environment on fusion frequency

While fusion is primarily influenced by genetic factors, non-genetic factors, such as abiotic conditions, may also play a role in the decision to fuse. Moreover, these conditions can fluctuate over time, such as during alternating periods of stress exposure, potentially altering the costs and benefits of fusion in different scenarios. To test this hypothesis, we examined fusion frequency in pairs of plasmodia composed of either clonal or fusion-compatible non-clonal plasmodia under conditions with and without salt. We found that salt significantly increased fusion frequency (𝜒^2^=39.787, *df*=1, *P*<0.001), with ∼50% fusion in the absence of salt compared to nearly 100% in its presence (Fig. 5). This effect was consistent in both pairs of clones and compatible non-clones. The genetic identity of the fusion partner (clonal versus compatible non-clonal) had no significant effect on fusion frequency (𝜒^2^=0.189, *df*=1, *P*=0.64).

**Figure 5.**
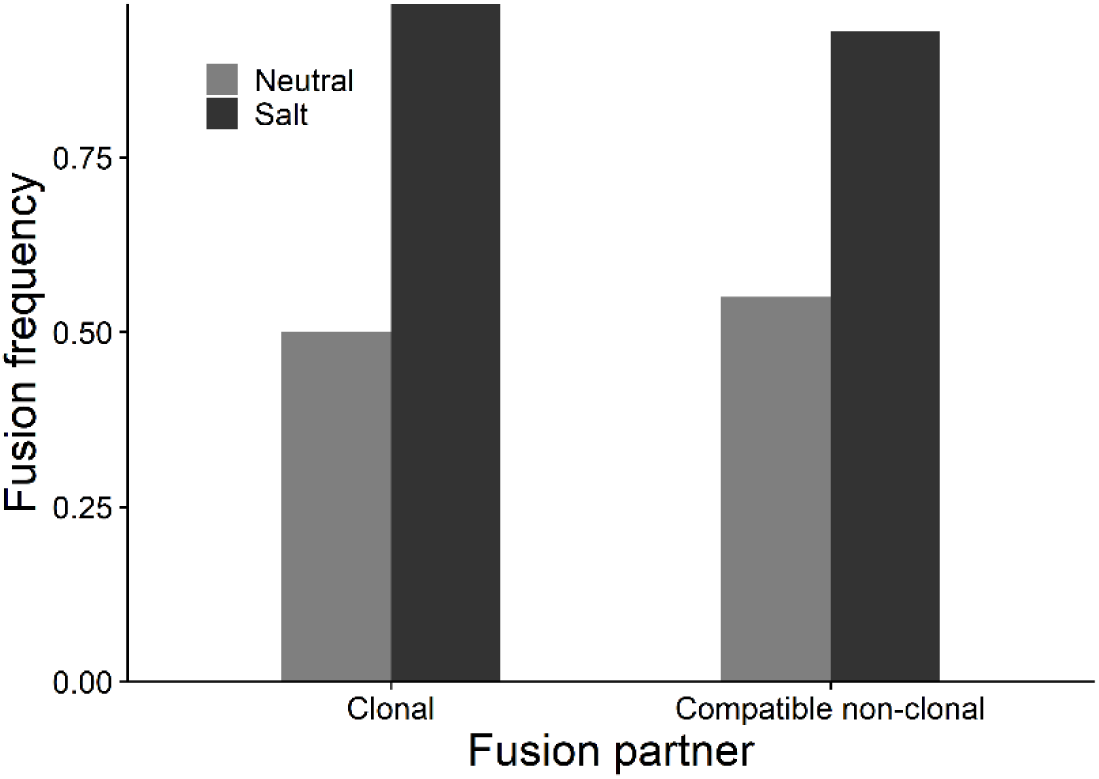
The influence of salt and fusion partner (clonal versus compatible non-clonal plasmodium) on fusion frequency.

### The benefits of increased size as a result of fusion

Next, we investigated whether the size-related benefit observed in larger plasmodia can be similarly achieved through fusion with another individual. This idea was initially tested using pairs of clonal plasmodia.

Comparing a single plasmodium to two fused clonal plasmodia we found that fused plasmodia crossed significantly wider salt compartments (Fig. 6A; 𝜒^2^=4.692, *df*=1, *P*=0.030). In addition, plasmodia that increased in size through fusion also crossed salt compartments with higher salt concentrations (Fig. 6B; 𝜒^2^=14.522, *df*=1, *P*<0.001).

**Figure 6.**
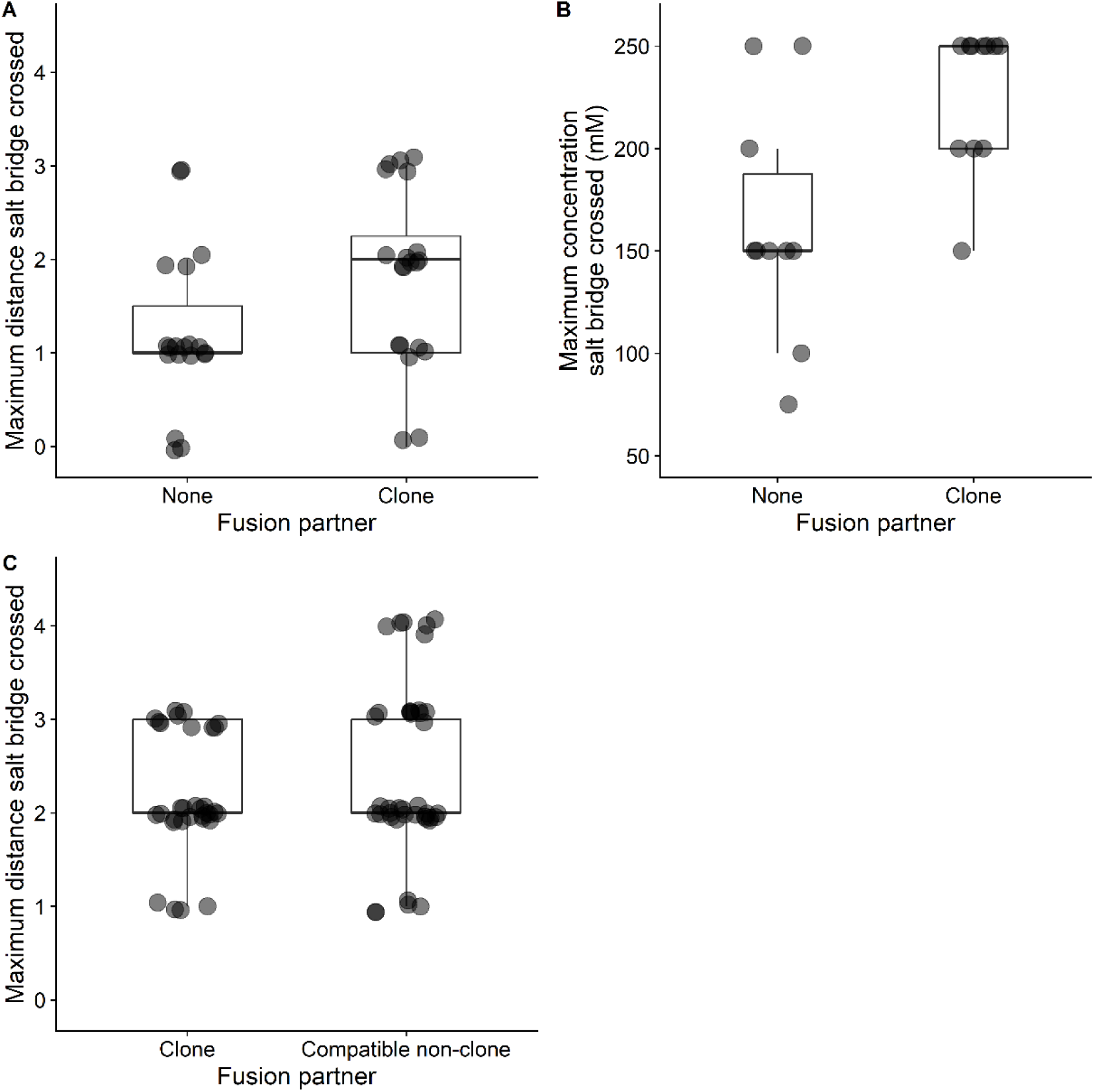
(A) The maximum distance of a 150mM salt compartment (in cm) crossed by a single network (fusion partner = none) versus a network made up of two clones. The initial size of the networks was the same, meaning that a fused network is ∼double the size of a single network. (B) The maximum concentration of the salt (mM) compartment crossed by a single network versus a network made up of two clones. (C) The maximum distance of a 150mM salt compartment (in cm) crossed by a network made up of two clones versus compatible non-clones.

Because we identified pairs of closely related plasmodia that successfully fused (Table 1), we could test whether size-related benefits differed between clonal and compatible non-clonal fusion partners. We found no significant difference in the maximum distance crossed between fused networks made up of clones and compatible non-clones (Fig. 6C; 𝜒^2^=1.617, *df*=1, *P*=0.20). Collectively, these results suggest that, at least in this set of tested pairs, fusion generally provides a size-related survival benefit, regardless of whether the fusion partner is an identical plasmodium or a compatible close relative.

## Discussion

Here we examined the frequency and consequences of plasmodial fusion between clonal and non-clonal strains of the slime mould *Physarum polycephalum*. We found a variety of fusion outcomes among close relatives, ranging from successful fusion to rejection, with or without a strong ‘aggressive’ reaction which seemingly impacted one or both individuals. In addition, we found that fusion frequency significantly increased when individuals were exposed to stress. These results confirm prior work that demonstrated that fusion is firstly determined by genetic factors (Poulter and Dee 1968; Carlile and Dee 1967), and additionally show that fusion frequency varies depending on abiotic conditions.

We observed variation in the fusion-rejection ratio among the three plasmodial lines tested, with fusion rates ranging from 3% to 63%. This variation likely reflects underlying differences in the homozygosity of fusion loci between different strains, where higher levels of homozygosity would increase the likelihood of successful fusion between siblings. Unfortunately, the molecular basis of fusion remains unknown, which precludes direct tests of this idea. Similarly, there is a complete lack of understanding of *P. polycephalum’s* genetic population structure (or its natural history), which limits our ability to assess the relevance of these differences in fusion frequencies to natural populations. Future studies should focus on co-isolating *P. polycephalum* lines to clarify how frequently encounters among close relatives and unrelated individuals prevail and the scale at which cooperation and competition occurs in nature.

We then tested the benefit of fusion-mediated size increase on survival. To do so, we first demonstrated that larger size confers the ability to cross wider salt bridges and bridges with higher salt concentrations. We next confirmed that similar size-benefits were obtained through fusion, even though fusion results in larger size much more rapidly than through growth. Interestingly, these benefits were obtained regardless of whether an individual fused with a clonal partner or a fusion-compatible close relative (sibling). Together these results indicate that, at least in the pairs of individuals tested here, the benefit of the size increase outweighs its potential costs associated from fusing with a non-clonal partner.

Prior work in other fusion-capable organisms has demonstrated that larger size benefits can result from enhanced resource acquisition, greater competitiveness in social interactions, and improved defence against predators and environmental stressors (Buss 1982, 1990; Feldgarden and Yund 1992; Stoner, Rinkevich, and Weissman 1999; Aanen et al. 2008; Bastiaans, Debets, and Aanen 2015). Similarly, we show that larger plasmodial networks exhibit greater resilience to salt stress in *Physarum*. One possibility is that this advantage stems from the dilution of the salt across a larger cytoplasmic volume, minimizing localized damage.

Additionally, larger networks may redistribute resources more effectively, repair damage more efficiently, and maintain higher nutrient reserves to counter stress, collectively enhancing the plasmodium’s ability to tolerate environmental challenges. At present, our results are restricted to a single environmental stressor. Although our findings add to our understanding of how size confers survival benefits in *Physarum*, further work is needed to determine the generality of these results in this and other systems.

Based on theory and empirical studies in other organisms that exhibit chimerism, we predicted that fused networks composed of genetically different individuals would show reduced network functioning—resulting from the possibility of inter-genotype conflicts—compared to those consisting of clones. For example, chimeric groups of the social amoebae *Dictyostelium discoideum* exhibit reduced migration relative to clonal groups of the same size, although chimerism in this case does not involve cell-cell fusion (Foster et al. 2002). Similarly, in the aggregative fruiting bacterium *Myxococcus xanthus*, within-group genotype richness showed a significant negative correlation with swarming rate and spore production (Mendes-Soares et al. 2014). Contrary to our expectations, however, we found similar outcomes in chimeric compared to clonal plasmodia. This may imply that, under the conditions tested, the genetic relatedness between lines in the chimeric networks and/or the size-related benefits of fusion are sufficiently high, and the potential costs for social exploitation are sufficiently low, for selection to promote cooperation. This idea is supported by our data showing that fusion frequencies increase after exposure to salt stress, suggesting that risks of fusion with non-clonal partners were smaller than the potential benefits obtained from increased size.

An important limitation of our results, and on conclusions regarding costs and benefits of fusion, is that our experiments only considered short-term effects of fusion (∼24 hours timeframe). It is possible is that costs associated with non-clonal fusion manifest over longer timescales. Earlier research, based on a limited number of lines, observed several outcomes: stable coexistence of both lines, competitive interactions leading to dominance of one line, or active elimination where one line eradicates the other (Carlile and Dee 1967; Lane and Carlile 1979; Schrauwen 1984). Among these, stable coexistence appears most plausible in our experiments, as significant competition within the chimeric plasmodium would likely have impaired functioning, resulting in reduced survival of chimeric compared to clonal plasmodia. Nonetheless, future work will need to explore longer-term post-fusion processes. This could include long-term studies of nuclear dynamics such as coexistence or competitive exclusion, using microscopy, as well as sequencing to quantify genetic composition in ‘older’ plasmodia and, importantly, spores. Together with elucidating the genetic structure of natural populations, this work will be crucial for understanding the evolutionary and ecological consequences of non-clonal fusion.

## Acknowledgments

The authors would like to thank Narie Sasaki, Shigeyuki Kawano, and Tetsuya Higashiyama for the *Physarum polycephalum* strains and Arjan Kortholt for the *Klebsiella pneumonia* strain. This work was funded by the Human Science Frontier Program (RGP0001/2021) to DR, K. Alim and M. Roper.

## Author contributions

CB and DR conceived the idea and designed the experiments, CB collected and analysed the data, CB and DR wrote the manuscript and approved the final version.

## Data accessibility

Data and scripts will be made available online after submission.

